# Revealing the secrets beneath grapevine and *Plasmopara viticola* early communication: a picture of host and pathogen proteomes

**DOI:** 10.1101/2021.12.27.474247

**Authors:** Joana Figueiredo, Rita B. Santos, Leonor Guerra-Guimarães, Céline C. Leclercq, Jenny Renaut, Lisete Sousa, Andreia Figueiredo

## Abstract

Plant apoplast is the first hub of plant-pathogen communication where pathogen effectors are recognized by plant defensive proteins and cell receptors and several signal transduction pathways are activated. As a result of this first contact, the host triggers a defence response that involves the modulation of several extra and intracellular proteins. In grapevine-pathogen interactions, little is known about the communication between cells and apoplast. Also, the role of apoplastic proteins in response to pathogens still remains a blackbox. In this study we focused on the first 6 hours after *Plasmopara viticola* inoculation to evaluate grapevine proteome modulation in the apoplastic fluid (APF) and whole leaf tissue. *Plasmopara viticola* proteome was also assessed enabling a deeper understanding of plant and pathogen communication. Our results showed that oomycete recognition, plant cell wall modifications, ROS signalling and disruption of oomycete structures are triggered in Regent after *P. viticola* inoculation. Our results highlight a strict relation between the apoplastic pathways modulated and the proteins identified in the whole leaf proteome. On the other hand, *P. viticola* proteins related to growth/morphogenesis and virulence mechanisms were the most predominant. This pioneer study highlights the early dynamics of extra and intracellular communication in grapevine defence activation that leads to the successful establishment of an incompatible interaction.

## 1. Introduction

As sessile organisms, plants have developed mechanisms to rapidly adapt to environmental changes and pathogen attack. To overcome pathogen challenges, plants must quickly recognize the invaders and mount a successful defence strategy. This chess game between the plant and pathogen is illustrated by the “zig zag model”, coined by Jones and Dangl in 2006 ^1^. In this model, the plants first recognize the pathogen-associated molecular patterns (PAMPs) through pattern recognition receptors (PRR) in the apoplast, leading to a PAMP-triggered immunity (PTI). The apoplast is extremely relevant for plant defence since is where plant and pathogen first meet and where recognition begins. Pathogen recognition culminates in the activation of plant defence responses including the induction of defence genes, production of reactive oxygen species (ROS) and deposition of callose. In a second phase, effector-triggered susceptibility or ETS, the pathogen overcomes the plant first response by deploying effectors that increase pathogen virulence, like Crinklers and RxLR effectors ^1^. In an incompatible interaction, the plant recognizes the pathogen effectors through R-proteins. The interaction between plant R-proteins and pathogen effectors results in an effector-triggered immunity (ETI), that ultimately results in a hypersensitive cell death response (HR) at the pathogen entry site ^1^. This interaction implies a tight communication between host and pathogen with the traffic of plant proteins and pathogen effector proteins between the apoplast and the intercellular space. While still misgraded, the study of plant apoplast is of extreme importance in plant-pathogen interactions so to identify proteins with a key role in plant defence strategies and better understand their interaction with pathogen molecules. Apoplast proteome was characterized for few plant models, constitutively or under abiotic/biotic stress, as for example, grapevine ^2,3^, poplar ^4^, tobacco ^5^, cowpea ^6^, rice ^7^, coffee ^8,9^ and Arabidopsis ^10^. However, few studies focus on uncovering apoplast proteome modulation considering plant-pathogen interactions and even less when considering woody crop plants, such as grapevine and obligatory biotrophic oomycetes, as the downy mildew etiological agent, *Plasmopara viticola*.

Grapevine (*Vitis vinifera* L.), is one of the major crops grown in temperate climates, however is highly susceptible to downy mildew, caused by *P. viticola* ((Berk. and Curt.) Berl. & de Toni) ^11^. In Europe, *P. viticola* infection leads to heavy crop losses and disease management for downy mildew relies on the massive use of pesticides in susceptible varieties in each growing season. This practice is against the demands of the European Union guidelines for pesticide reduction and sustainable viticulture (Directive 2009/128/EC), so the search for more plant- and environment-friendly solutions is imperative.

In a modern viticulture context, the development of grapevine crossing lines, in breeding programs, is a very well established and accepted strategy to fight against the excessive use of pesticides. These crossing lines are the result of the introgression of pathogen resistant genes, present in Asian and American *Vitis* species, with genes related to the good quality of grapes for wine production, present in susceptible grapevine cultivars. The result is a cultivar that present desired characteristics for wine producers at the same time that resists more to pathogen attack. ‘Regent’ is a successful example of breeding for resistance and harbours RPV3.1 resistance to *P. viticola* loci ^12^. Several studies have been performed in ‘Regent’-*P. viticola* interaction with the aim to better understand the molecular mechanisms and the key molecules that are responsible to the well-known tolerance that this crossing line has against *P. viticola* ^13–17^.

In a climate change scenario, viticulture will face new emerging diseases as well as several outbreaks of the established diseases, such as downy mildew. Thus, a comprehensive knowledge on the grapevine strategies to overcome pathogens, mainly in cultivars with some resistance level, as well as the evolution of pathogen infection mechanisms is paramount to tackle this challenge. Thus, in the present study, we have focused on the early communication between grapevine and *P. viticola* and assessed, for the first time, grapevine apoplast proteome modulation and *P. viticola* proteome and secretomes. We have focused on the first 6 hours post inoculation (hpi), as the events occurring in this time-point were previously shown to be crucial for the outcome of the interaction. We have also highlighted extra and intra-cellular communication pathways by comparing the proteome modulation in the apoplast and in the whole leaf. Up to our knowledge this is the first time where grapevine apoplast and whole leaf proteome communication is revealed during host-pathogen interaction and also the first *P. viticola* proteome sequencing. Our results elucidate the interaction between grapevine and *P. viticola* proteins taking place in the apoplast and how the plant and pathogen proteomes evolve at the first stages of infection.

## 2. Results

### 2.1 Early ‘Regent’ APF proteome modulation under *P. viticola* infection

The impact of *P. viticola* infection in the modulation of APF proteome in ‘Regent’ leaves was analysed at 6 hours post inoculation (hpi). By a principal components analysis (PCA), a clear distinction between the proteome of inoculated and mock-inoculated (control) samples was obtained (Fig.1). The distribution of the biological replicates within the PCA scores plot indicates the absence of unwanted variation in the dataset, increasing the confidence in the reproducibility of the differential accumulation analysis.

**Fig.1.**
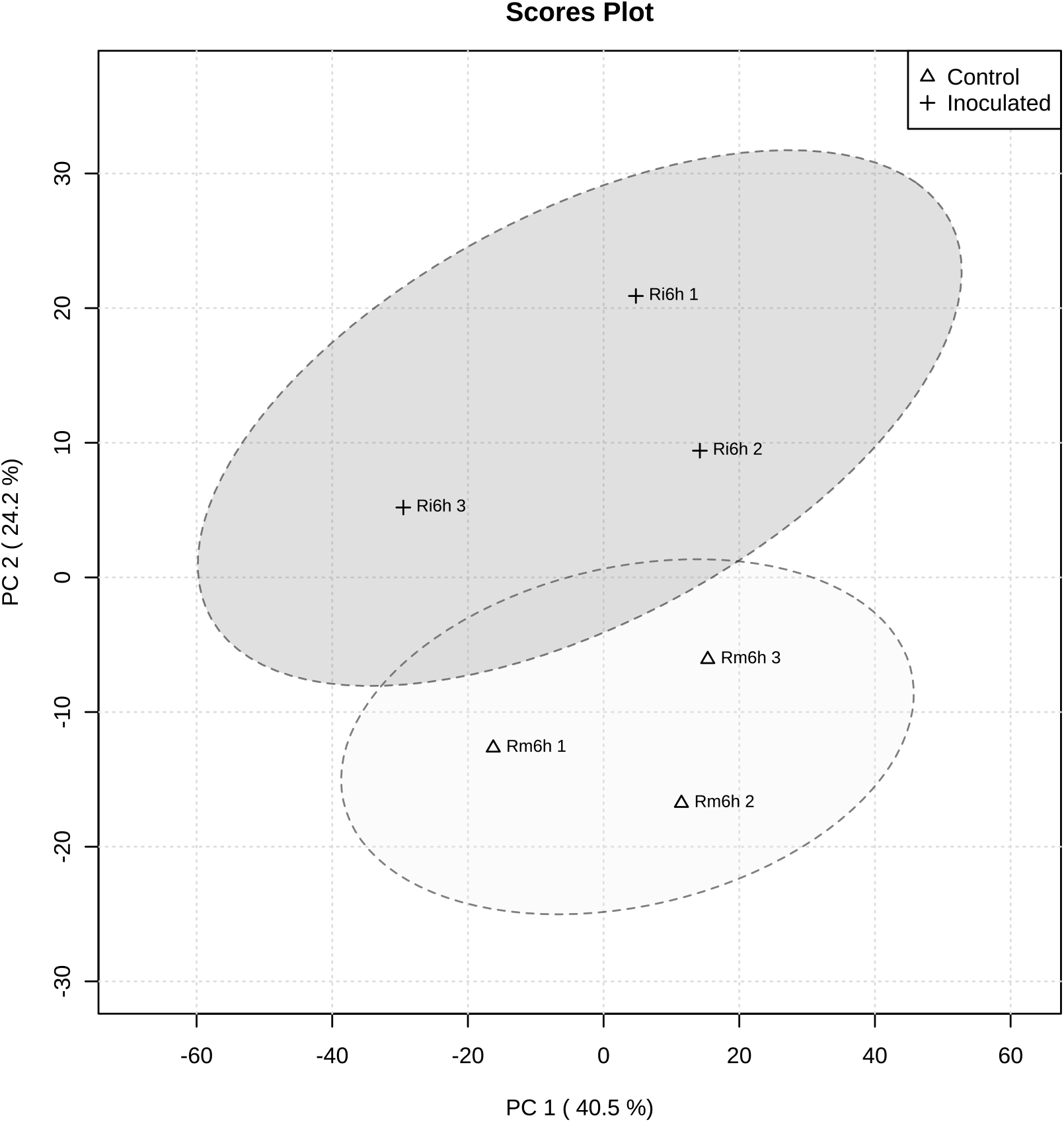
Principal component analysis of the differential protein profiles in *V. vinifera* cv. ‘Regent’ at 6 hours post-inoculation with *P. viticola*. The plot shows principal component 1 (PC1) on X axis and principal component 2 (PC2) on Y axis, together they explain 64.7% of protein abundance variability. Ri: ‘Regent’ inoculated samples; Rm: ‘Regent’ mock-inoculated samples.

When comparing APF inoculated samples with mock-inoculated samples, a hundred and eighteen proteins were differentially accumulated (DAPs; 74 up accumulated and 44 down accumulated). These proteins are mainly related to stress response, signal transduction, cell wall metabolism, transport and protein metabolism (Fig.2). At 6hpi, *P. viticola* infection leads to an increase in the presence of plant stress response proteins, like glucan endo-1,3-β-glucosidases, disease resistance proteins RPV1-like and RUN1-like [associated to Resistance to *P. viticola* (RPV) loci and Resistance to *Uncinula necator* (RUN) loci], receptor-like protein kinase FERONIA and GDSL esterases/lipases. An up accumulation of LRR kinases related to signal transduction occurs. ROS-related proteins, like peroxidase 4-like and xanthine dehydrogenase 1-like isoform X1, were also detected in ‘Regent’ APF after oomycete challenged.

**Fig.2.**
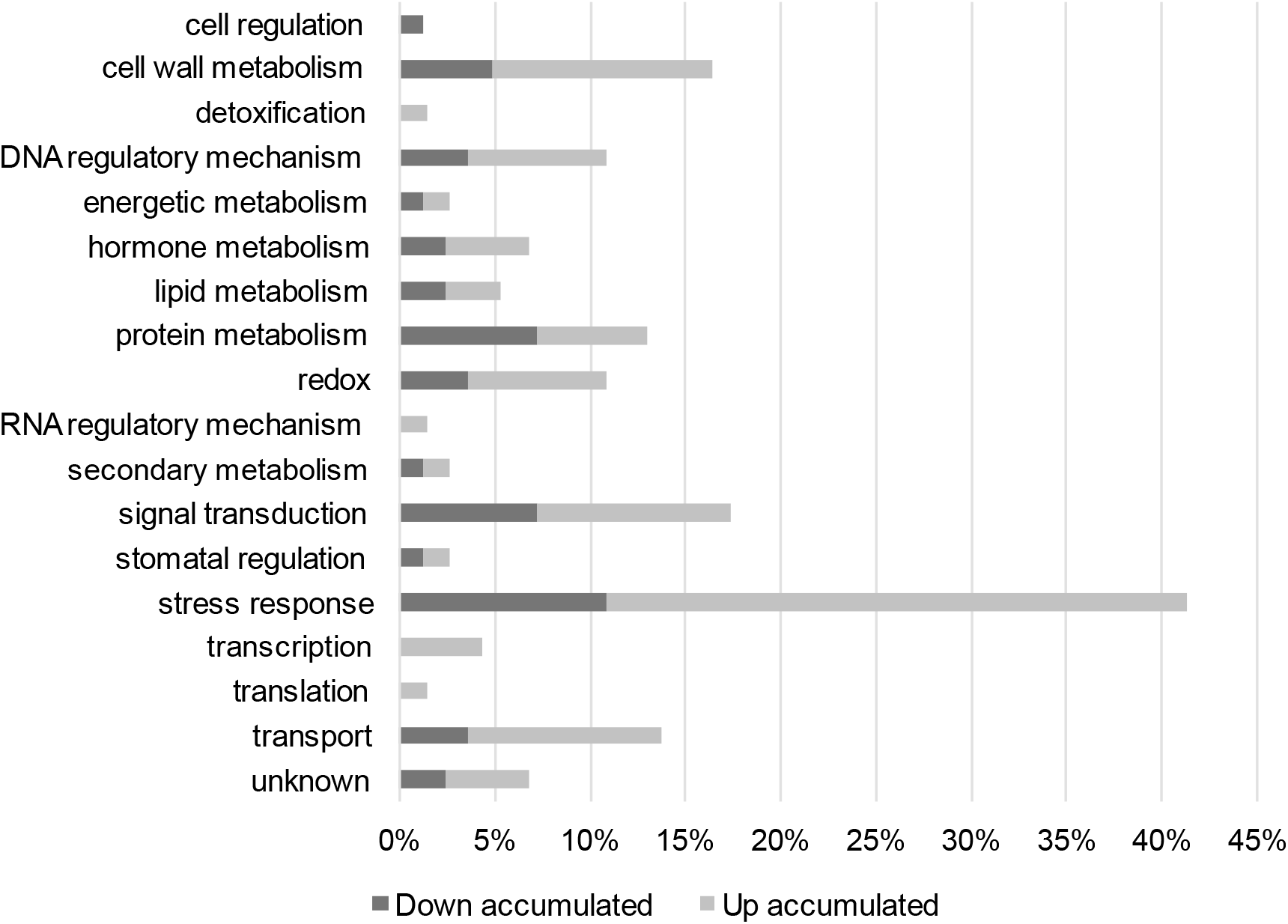
Biological process annotation of the differentially accumulated ‘Regent’ APF proteins, 6 hours post inoculation with *P. viticola*. Dark grey bars: percentage of down accumulated proteins; Light grey bars: percentage of up accumulated proteins.

Several proteins involved in ‘Regent’ defence mechanism were modulated at 6hpi (Table 1). Indeed, apoplastic proteins related to oomycete perception, that may lead to the activation of several defence signalling pathways, were found to be accumulated (Table 1). Apoplastic proteins associated with the remodelling of plant cell wall were also identified as well as proteins associated with auxin signalling and its regulation in response to *P. viticola* infection (Table 1). Moreover, several proteins associated to ROS production and signalling were modulated at this early time-point of infection (Table 1). Lastly, plant proteins involved in the disruption of oomycete structures were also identified (Table 1).

**Table 1.** ‘Regent’ APF proteins up accumulated at 6hpi with *P. viticola*, involved in key cellular pathways.

| NCBI accession | NCBI description | Biological process | log2(FC) |
| --- | --- | --- | --- |
| **Modulation of plant physical barriers** |  |  |  |
| XP_002281842.1 | ABC transporter G family member 32 (ABCG32) | stress response | 19,00 |
| XP_002263127.1 | fatty acyl-CoA reductase 3-like (FAR3) | lipid metabolism | 18,83 |
| NP_001268091.1 | pectinesterase/pectinesterase inhibitor PPE8B-like (PPE8B) | cell wall modification | 15,93 |
| XP_002277293.4 | pectinesterase | cell wall metabolism | -1,80 |
| **Modulation of plant plasma membrane proteins in response to infection** |  |  |  |
| XP_019071787.1 | disease resistance protein RPV1-like isoform X1 (RPV1) | stress response | 19,12 |
| XP_019073586.1 | disease resistance protein RUN1-like isoform X1 (RUN1) | stress response | 17,99 |
| XP_010644327.1 | disease resistance protein At4g27190-like | stress response | 4,69 |
| XP_002280315.3 | probable disease resistance protein At1g61300 | stress response | 18,99 |
| XP_019077695.1 | probable disease resistance protein At1g61300 | stress response | 4,47 |
| XP_010654733.1 | probable disease resistance protein At5g63020 | stress response | 21,94 |
| XP_010645387.1 | TMV resistance protein N | stress response | 21,70 |
| XP_019078946.1 | TMV resistance protein N | stress response | 17,22 |
| XP_019075299.1 | leucine-rich repeat receptor protein kinase MSP1-like | signal transduction | 16,43 |
| XP_002267269.1 | probable leucine-rich repeat receptor-like protein kinase At1g35710 | signal transduction | 16,39 |
| XP_002282474.2 | serine-threonine protein kinase, plant-type, putative | signal transduction | 18,49 |
| XP_010660578.1 | receptor-like protein kinase FERONIA (FERONIA) | stress response | 18,38 |
| XP_010664467.1 | ATPase 9, plasma membrane-type | transport | 19,39 |
| XP_002279498.1 | putative calcium-transporting ATPase 13, plasma membrane-type (Ca<sup>2+</sup>-ATPase) | transport | 15,64 |
| XP_002283826.1 | protein unc-13 homolog | stomatal regulation | 18,55 |
| XP_010647591.2 | ankyrin repeat-containing protein ITN1-like isoform X6 (ITN1) | stress response | 16,63 |
| **Activation of auxin signalling** |  |  |  |
| XP_002274153.1 | protein kinase PINOID | hormone signalling | 22,45 |
| XP_002277611.1 | protein NDL2 | hormone signalling | 16,72 |
| XP_002265864.1 | ubiquitin-NEDD8-like protein RUB2 | protein metabolism | 16,62 |
| **Regulation of ROS during plant defence response** |  |  |  |
| XP_002269918.1 | peroxidase 4-like | redox | 21,05 |
| XP_002274392.1 | purple acid phosphatase (PAP) | energetic metabolism | 20,55 |
| XP_002285473.1 | xanthine dehydrogenase 1-like isoform X1 (XDH1) | redox | 16,51 |
| **Disruption of oomycete structures** |  |  |  |
| XP_010664681.1 | glucan endo-1,3-beta-D-glucosidase | stress response | 19,85 |
| XP_002283647.1 | glucan endo-1,3-beta-glucosidase | stress response | 15,09 |
| XP_002268991.2 | GDSL esterase/lipase (GELP) | stress response | 15,72 |
| XP_002276525.1 | GDSL esterase/lipase At4g01130 (GELP) | stress response | 6,00 |

### 2.2 ‘Regent’ whole leaf proteome modulation at 6h post *P. viticola* infection

Whole leaf proteome of ‘Regent’ during *P. viticola* infection time-course was obtained in 2020 ^13^. Raw data deposited on the Pride database of the 6hpi was re-analysed following the same pipeline that was implemented for the APF proteome analysis. Hundred and fifty-two DAPs were identified. Of those, 69 proteins were up accumulated and 83 were down accumulated. These proteins were mainly related to stress response, energy and secondary metabolisms and translation (Fig.3). At 6hpi, *P. viticola* infection induces a modulation in the abundance of several stress-related proteins, like heat shock proteins, cysteine proteinases and glucanases. In addition, a great number of photosynthesis-related proteins are down accumulated in ‘Regent’ leaves in response to infection. Translation and signal transduction-related proteins, like ribosomal proteins and serine/threonine protein kinase (RSTK), respectively, were up accumulated after *P. viticola* infection.

**Fig.3.**
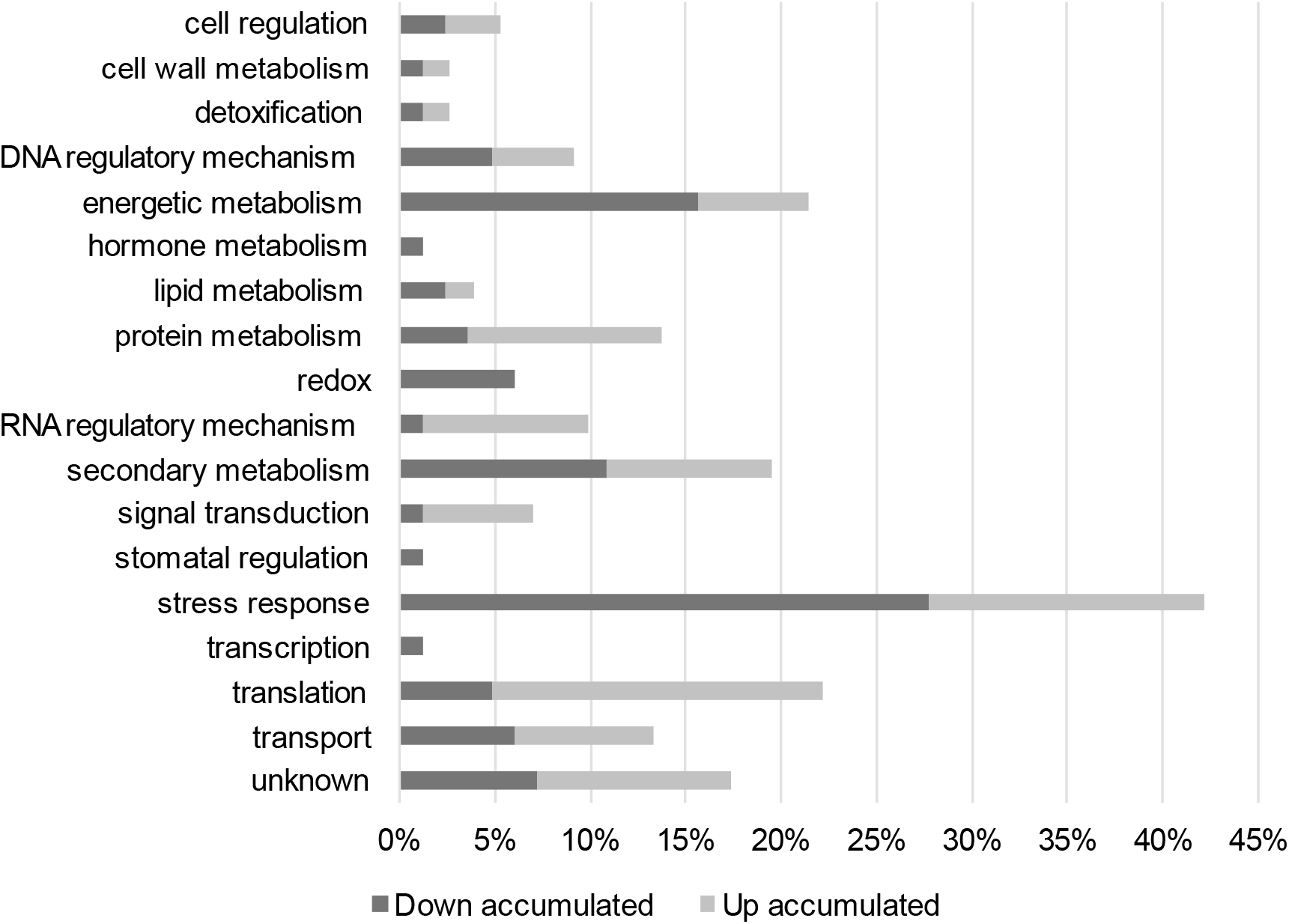
Biological process annotation of the differentially accumulated ‘Regent’ whole leaf proteins, 6 hours post inoculation with *P. viticola*. Dark grey bars: percentage of down accumulated proteins; Light grey bars: percentage of up accumulated proteins.

The functional annotation of several of the proteins identified in whole leaf are closely related with the pathways that were found to be modulated in APF proteins (Table 2). Indeed, we have identified whole leaf proteins involved in the modulation of plant physical barriers and activation of plant defence signalling through plasma membrane receptors, regulation of ROS levels, and disruption of oomycete structures (Table 2). Also, proteins associated with calcium signalling, and intracellular trafficking vesicles were modulated in the whole leaf context (Table 2).

**Table 2.** Regent’ whole proteins up accumulated at 6hpi with *P. viticola*, involved in key cellular pathways.

| NCBI accession | NCBI description | Biological process | log<sub>2</sub>(FC) |
| --- | --- | --- | --- |
| **Modulation of plant physical barriers and plasma membrane receptors in response to infection** |  |  |  |
| CAN83165.1 | pectinesterase inhibitor 9 (PMEI9) | cell wall metabolism | 2,39 |
| CBI32865.3 | alpha-L-arabinofuranosidase 1 (ASD1) | cell wall metabolism | -18,97 |
| XP_002267434.3 | serine/threonine protein kinase (RSTK) | signal transduction | 6,90 |
| XP_002265462.3 | serine/threonine-protein kinase pakA (RSTK) | signal transduction | 17,31 |
| **Regulation of ROS during plant defence response** |  |  |  |
| CBI31928.3 | peroxisome biogenesis protein 19-2-like (PEX19-2) | protein metabolism | 19,28 |
| CBI32544.3 | protein TIC 62, chloroplastic isoform X1 | transport | 18,74 |
| CAN66554.1 | succinate dehydrogenase assembly factor 4, mitochondrial (SDHAF4) | energetic metabolism | 17,63 |
| XP_002283860.1 | 15.7 kDa heat shock protein, peroxisomal | stress response | 20,89 |
| CAN67665.1 | 17.3 kDa class II heat shock protein-like | stress response | 6,61 |
| XP_003634522.1 | peroxidase 12 | redox | -21,36 |
| XP_002285652.2 | peroxidase A2-like | redox | -8,08 |
| XP_010647098.1 | polyphenol oxidase, chloroplastic-like | redox | -20,80 |
| CBM39273.1 | 18.2 kDa class I heat shock protein | stress response | -20,66 |
| CBI30632.3 | 28 kDa heat- and acid-stable phosphoprotein | stress response | -19,75 |
| CBM39216.1 | class I heat shock protein | stress response | -21,56 |
| CBI23075.3 | small heat shock protein, chloroplastic | stress response | -21,87 |
| **Disruption of oomycete structures** |  |  |  |
| CBI32343.3 | endo-1,3;1,4-beta-D-glucanase-like | stress response | -21,33 |
| CBI26171.3 | endo-1,3;1,4-beta-D-glucanase-like isoform X3 | stress response | -19,29 |
| CAN71820.1 | glucan endo-1,3-beta-glucosidase | stress response | -20,49 |
| CBI36040.3 | profilin 1 | cell regulation | 20,96 |
| **Modulation of calcium signalling** |  |  |  |
| CBI15387.3 | calcium sensing receptor, chloroplastic | stomatal regulation | -19,74 |
| CBI24493.3 | CDGSH iron-sulfur domain-containing protein NEET | secondary metabolism | -7,88 |
| **Increase of protein trafficking** |  |  |  |
| CBI28256.3 | putative clathrin assembly protein At5g35200 | stress response | 21,36 |

### 2.3 *P. viticola* proteome during infection establishment

We have sequenced for the first time the *P. viticola* proteome, obtained from grapevine leaves apoplast at 6hpi. Sixty proteins were identified being mainly involved in two biological processes: growth/morphogenesis (e.g. cell division cycle 5 and β-glucan synthesis-associated SKN1) and virulence (RxLR proteins and serine protease trypsin’s). Proteins involved in signalling processes like agc kinase (ACG), serine threonine kinase and small GTP-binding Rab28 were also identified (Table 3).

**Table 3.** *P. viticola* proteins, identified in ‘Regent’ apoplast after 6hpi, involved in virulence and growth/morphogenesis mechanisms.

| Protein name | Protein code (INRA Database) |
| --- | --- |
| **Proteins involved in virulence mechanisms** |  |
| Coproporphyrinogen III oxidase (CPOX) | PVIT_0003600.T1 |
| RxLR-like protein (RxLR) | PVIT_0014146.T1 |
| RxLR-like protein (RxLR) | PVIT_0014142.T1 |
| Serine protease trypsin | PVIT_0011817.T1 |
| Serine protease trypsin | PVIT_0011837.T1 |
| Serine threonine kinase | PVIT_0018302.T1 |
| Tetratricopeptide repeat 26 | PVIT_0013696.T1 |
| TKL kinase (TKL) | PVIT_0016228.T1 |
| **Proteins involved in growth/morphogenesis mechanisms** |  |
| Beta-glucan synthesis-associated SKN1 (SKN1) | PVIT_0022780.T1 |
| CAMKK kinase (CaMK) | PVIT_0013015.T1 |
| Cell division cycle 5 (CDC5) | PVIT_0005546.T1 |
| CMGC CDK kinase (CDK) | PVIT_0011642.T1 |
| CMGC MAPK kinase (MAPK) | PVIT_0009065.T1 |
| FAD synthase-like | PVIT_0001050.T1 |
| Serine threonine kinase | PVIT_0018302.T1 |
| **Proteins involved in both mechanisms** |  |
| AGC kinase (ACG) | PVIT_0009891.T1 |
| Calpain-like protease | PVIT_0019872.T1 |
| Small GTP-binding Rab28 | PVIT_0005551.T1 |

## 3. Discussion

### 3.1 *P. viticola* leads to a broad modulation of ‘Regent’ APF and whole leaf proteomes

During grapevine-*P viticola* interaction, the apoplast compartment is the first hub where plant and pathogen secretomes meet. Several proteins are crucial for the outcome of the interaction, both from the host or pathogen sides. In the apoplast, processes involving pathogen recognition through membrane receptors that activate signal transduction pathways for expression of host defence-associated genes or proteins that directly communicate with pathogen molecules inhibiting infection progress are essential. Considering the whole leaf tissue, trafficking of several proteins to respond to the plant defence requirements must be activated as well as processes that lead to a broad activation of defence-related mechanisms. Thus, communication between the apoplast and the host intracellular organelles is essential for a concerted and quick defence response against the pathogen. Moreover, during the interaction, *P. viticola* develops its infection structures, namely hyphae culminating in plant cell invasion and development of haustorium for feeding. In the first hours of interaction, host and pathogen communications are expected to be very dynamic and to define the outcome of the interaction.

#### 3.1.1 The dual battle at the gate: host strengthens its physical barriers while the pathogen triggers plant cell wall degradation

The cuticle is a barrier coating the outer surface of epidermal cells of organs of the aerial parts of the plants. It protects against water loss and various abiotic and biotic stresses ^18^. In ‘Regent’ APF, two cuticle-related proteins were found to be up accumulated after *P. viticola* infection, the ABC transporter G family member 32 (ABCG32) and the fatty acyl-CoA reductase 3-like protein (FAR3), (Table 1; Fig.4). These ABC transporters have been frequently shown to be involved in pathogen response, surface lipid deposition and transport of the phytohormones auxin and abscisic acid ^19,20^. In Arabidopsis, the ABCG32 was reported to be involved in cuticle formation, most likely by exporting cutin precursors from the epidermal cell ^21^. The fatty acyl-CoA reductase 3-like protein is involved in cuticular wax biosynthesis ^22^. In incompatible grapevine-*P. viticola* interaction, such as the one that occurs in ‘Regent’, a higher abundance of these proteins leads to the hypothesis that the host activates processes to promote the strengthening the cuticular barrier in order restrain pathogen penetration (Fig.4).

**Fig.4.**
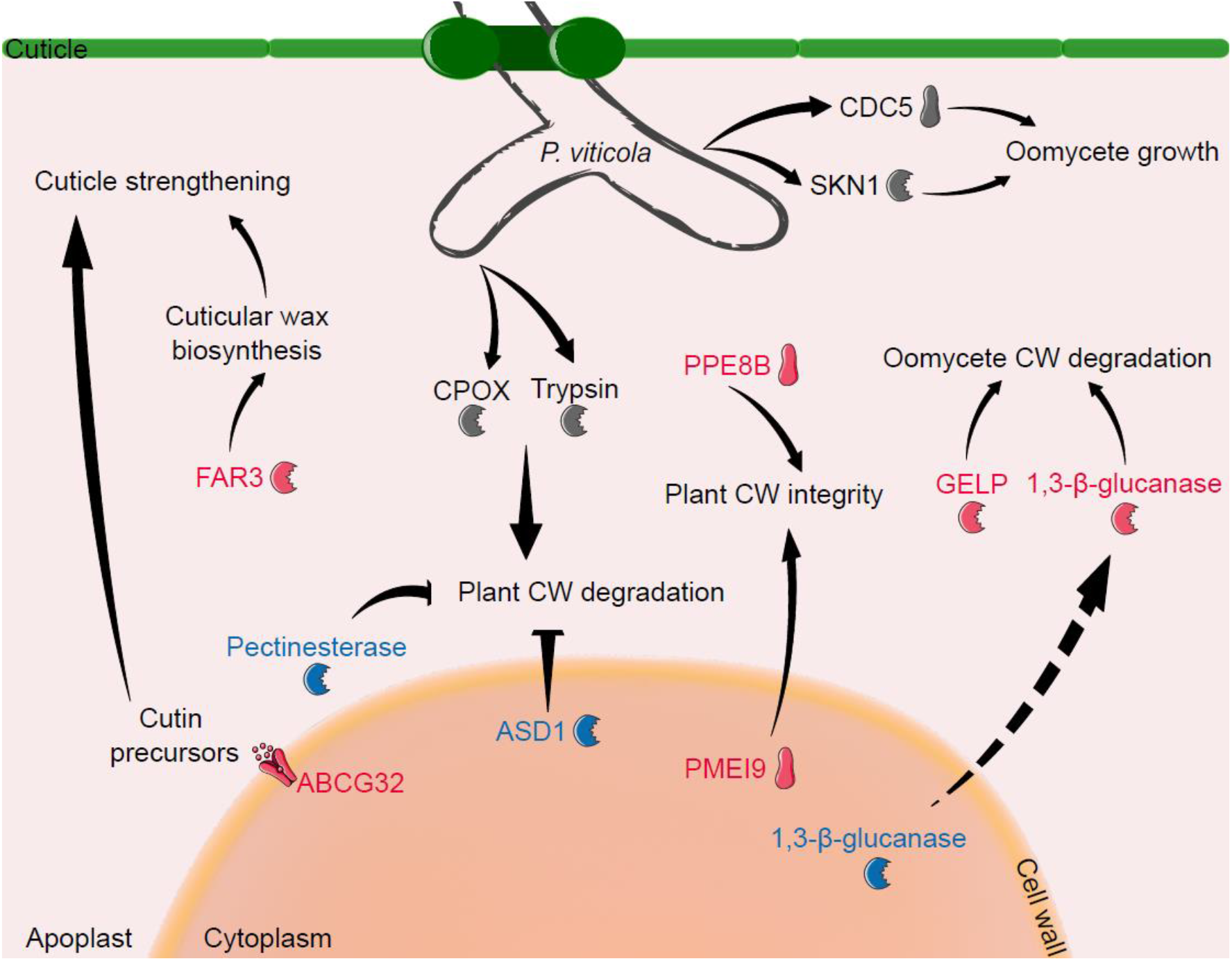
Triggering of host cell wall degradation by *P. viticola* through CPOX and trypsin proteins secretion while in the host several proteins associated to the strengthening of the physical barriers (cuticle – eg ABCG32, FAR3; cell wall - eg PMEI9, PPE8B) are positively modulated. At the host also a negative modulation of proteins involved in cell wall degradation is promoted, moreover, proteins as 1.3-β-glucanases are translocated to the APF for pathogen cell wall degradation. Proteins represented in red are positively modulated, proteins represented in blue are negatively modulated (compared to mock-inoculated control). Pathogen proteins are represented in grey. 1,3-β-glucanase – includes endo-1,3;1,4-beta-D-glucanase-like, endo-1,3;1,4-beta-D-glucanase-like isoform X3 and glucan endo-1,3-beta-glucosidase; ABCG32 - ABC transporter G family member 32; ASD1 - alpha-L-arabinofuranosidase 1; CDC5 - cell division cycle 5; CPOX - coproporphyrinogen III oxidase; CW - cell wall; GELP - GDSL esterase/lipase; FAR3 - fatty acyl-CoA reductase 3-like protein; PMEI9 - pectinesterase inhibitor 9; PPE8B - pectinesterase/pectinesterase inhibitor PPE8B-like; SKN1 - beta-glucan synthesis-associated SKN1.

Also, plants have developed a system for sensing pathogens and monitoring the cell wall integrity, upon which they activate defence responses that lead to a dynamic cell wall remodelling required to prevent pathogen progression. Plant cell wall-associated proteins were found to be differentially modulated in ‘Regent’ APF in response to *P. viticola*. A down accumulation of a pectinesterase, a protein involved in plant cell wall degradation as well as an up accumulation of pectinesterase/pectinesterase inhibitor PPE8B-like protein (PPE8B), (Table 1; Fig.4). This protein is involved in plant cell wall reconstruction and, in cotton, genes encoding this type of protein are specifically up regulated in plant resistant variety upon *Aspergillus tubingensis* infection ^23^.

In the whole leaf proteome of ‘Regent’, the abundance of pectinesterase inhibitor 9 (PMEI9) increased, a protein involved in resistance to pathogens ^24^, and a decrease in the accumulation of alpha-L-arabinofuranosidase 1, a protein involved in cell wall degradation ^25^, was detected (Table 2; Fig.4). The whole leaf and apoplast proteomes are modulated as a defence strategy to prevent cell wall degradation, maintaining its integrity and thus inhibiting the entry of the oomycete. On the other hand, proteins involved in plant cell wall degradation were found in *P. viticola* proteome (Table 3; Fig.4), namely: coproporphyrinogen III oxidase, which is a peroxidase with the ability to degrade lignin, one of the components of plant cell walls ^26^; trypsins, a serine proteases family, also identified in the secretomes of several fungus ^27^, and linked to pathogenicity against plant hosts ^28^. In *P. viticola*, the two identified trypsins were predicted to be apoplastic effectors and so it is expected a direct interaction with plant molecules for cell wall degradation.

During plant-pathogen interactions, plant cell wall is a dynamic structure that functions as a barrier that pathogens need to breach to colonize the plant tissue. Biotrophic pathogens, like *P. viticola*, require a localized and controlled degradation of the cell wall to keep the host cells alive during feeding. The regulation of the abundance of these cell wall-related proteins in the apoplast and whole leaf of ‘Regent’ suggests an adaptation of the grapevine proteome to prevent cell wall disruption while *P. viticola* secrets proteins that degrade de cell wall to invade the plant cell.

#### 3.1.2 *P. viticola* recognition and signalling events are established as soon as 6hpi

The first layer of plant defence relies on the recognition of conserved microbe-associated molecular patterns (MAMPs) by the so-called pattern recognition receptors (PRRs). PRRs are generally plasma membrane receptors which are often coupled to intracellular kinase domains ^29^. The second layer of plant immunity depends on the ability of the plant to recognize the pathogen effectors, like RxLRs, by disease resistance proteins (R) and trigger a robust resistance response ^30^. Several states such as oxidative burst, cell wall strengthening, induction of defence gene expression, and rapid cell death at the site of infection (hypersensitive response) occur in downstream cellular events leading to the establishment of an incompatible interaction ^31^. These local hypersensitive responses can trigger long-lasting systemic responses (systemic acquired resistance (SAR)) that prime the plant for resistance against a broad spectrum of pathogens ^32,33^.

We have identified several virulence-related proteins that are secreted by *P. viticola* or that are present in oomycete infection structures (Table 3; Fig.5). RxLR effectors were detected in *P. viticola* proteome as soon as 6hpi and were predicted to be secreted to the apoplast. These proteins are key players in virulence for downy mildew species ^34^ since they are known to defeat plant immune responses through many routes, which include reprogramming host gene expression, altering RNA metabolism, and binding to host proteins involved in signalling ^35^. On the host side, eight R-proteins were up-accumulated in the APF after *P. viticola* infection (Table 1; Fig.5), including RPV1-like isoform X1 and RUN1-like isoform X1, which confer resistance to multiple downy and powdery mildews, respectively, by promoting cell death ^36–38^. Also, an up accumulation of several RSTK was observed both in APF (Table 1) and in whole leaf proteome (Table 2). These receptors are involved in a wide array of processes ranging from developmental regulation to disease resistance, including activation of signal transduction for plant defence response initiation ^39,40^. One of the identified receptors in APF is the FERONIA (Fig.5). FERONIA is a plant recognition receptor kinase which plays a significant role in plant immune system. In *Catharanthus roseus*, FERONIA acts as a sensor of cell wall integrity challenged by the host-pathogen interaction and further triggers downstream immune responses in the host cell. In response to changes in the cell wall, FERONIA induces the release of ROS (reviewed in ^41^). Furthermore, in spinach response to infection with the oomycete *Peronospora effusa*, an up-regulation of the genes encoding for the FERONIA and a LRR receptor-like RSTK was observed ^42^. These results highlight the importance of these receptors in the plant cell membrane and in the first line of defence in plant strategies to stop oomycete proliferation.

**Fig.5.**
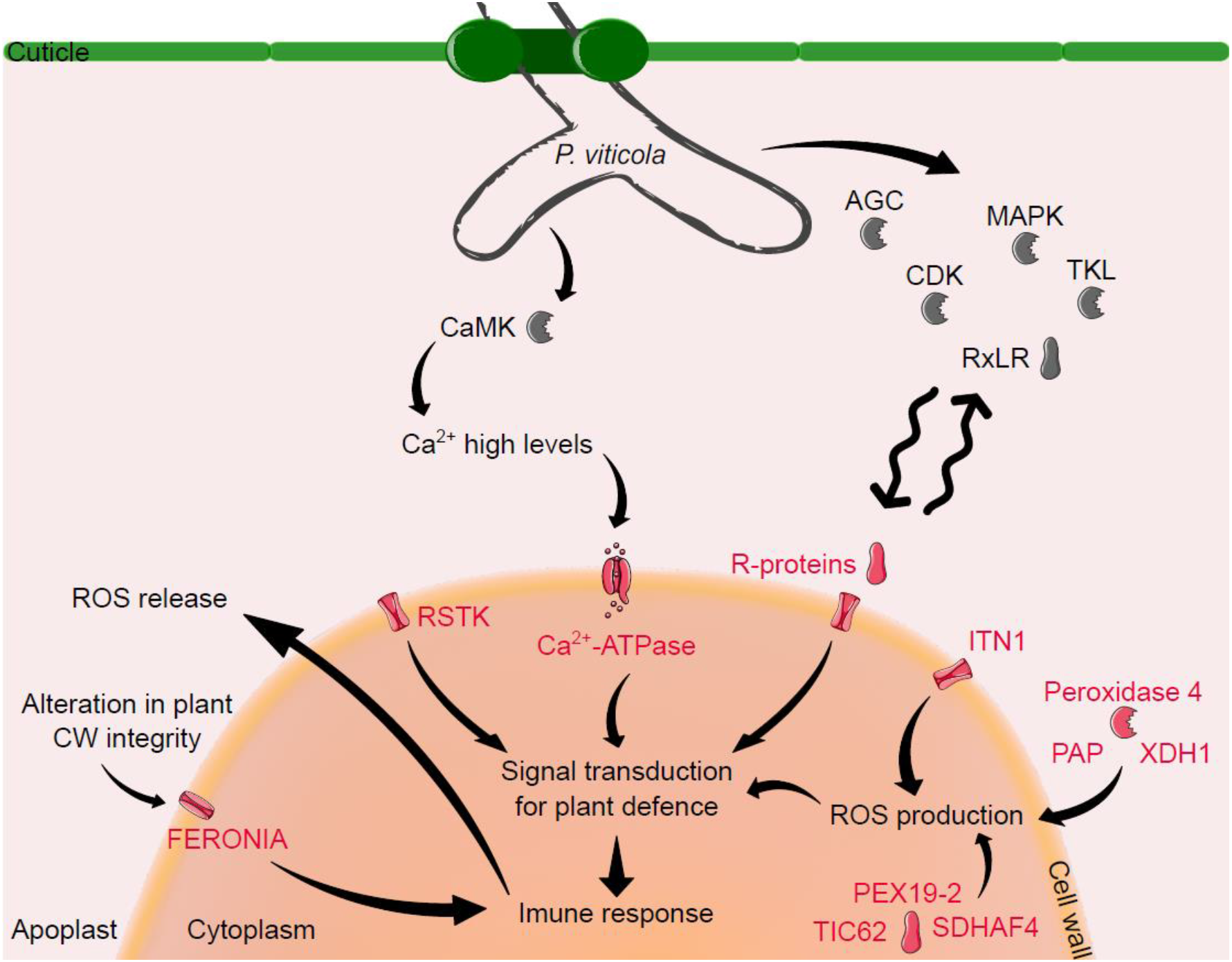
First communication events between host and pathogen: pathogen effectors are recognized by host R proteins leading to a broader remodulation of proteins related to ROS production, signal transduction and immune response establishment. Also, *P. viticola* is able to perceive host molecules activating signalling transduction pathways. Proteins represented in red are positively modulated, proteins represented in blue are negatively modulated (compared to mock-inoculated control). Pathogen proteins are represented in grey. AGC - AGC kinase; Ca^2+^-ATPase - putative calcium-transporting ATPase 13; CaMK - CAMKK kinase; CDK - CMGC CDK kinase; CW - cell wall; FERONIA - receptor-like protein kinase FERONIA; ITN1 - ankyrin repeat-containing protein ITN1-like isoform X6; MAPK - CMGC MAPK kinase; PAP - purple acid phosphatase; PEX19-2 - peroxisome biogenesis protein 19-2-like; RSTK - serine/threonine protein kinase; ROS – reactive oxygen species; RxLR - RxLR-like proteins; R-proteins – includes RPV1-like isoform X1 and RUN1-like isoform X1 and RSTKs; SDHAF4 - succinate dehydrogenase assembly factor 4; TIC62 - protein TIC 62; TKL - TKL kinase; XDH1 - xanthine dehydrogenase 1.

Moreover, an up accumulation of a H^+^-ATPase (ATPase 9) was also detected in ‘Regent’ APF at 6hpi. Plasma membrane H^+^-ATPases maintain low cytoplasmic concentrations of H^+^ and are dynamically regulated by biotic and abiotic events, being involved in signalling mechanisms during plant-pathogen interaction (reviewed in ^43^). In response to internal and/or external cues, the trafficking of these plant pumps, to and from the plasma membrane, is also highly regulated. Indeed, we have identified an up accumulation of a protein involved in H^+^-ATPases translocation for the plasma membrane, the protein unc-13 homolog (Table 1). In Arabidopsis, a protein unc-13 homolog (PATROL1) controls the translocation of a H^+^-ATPase to the plasma membrane during stomata opening ^44^. PATROL1 resides in the endosome and moves to and from the plasma membrane in response to environmental stimuli. When the stomata open, intracellular vesicles incorporate the plasma membrane and PATROL1 may be carried with them to tether the H^+^-ATPase into the plasma membrane ^44^. Modifications of the H^+^-ATPases concentration in the plasma membrane act as an alternative mean to control stomata opening ^45^ and in the case of grapevine-*P. viticola* interaction, the stomata are the entry sites of the oomycete. Despite the role of these ATPases in grapevine remains to be elucidated, an up accumulation of ATPase 9 and protein unc-13 homolog could suggest its involvement in stomata opening regulation during ‘Regent’-*P. viticola* interaction.

During ‘Regent’ response to *P. viticola* infection, the up accumulation of another transmembrane protein (TMP), ankyrin repeat-containing protein ITN1-like isoform X6 (ITN1), was also detected (Table 1; Fig.5). Plant TMPs are essential for normal cellular homeostasis, nutrient exchange, and responses to environmental cues ^46^. ITN1 has been implicated in diverse cellular processes such as signal transduction and, in Arabidopsis, this protein was proposed to be related with abscisic acid signalling pathway and promotion of ROS production ^47^.

#### 3.1.3 Auxin signalling plays an important role in apoplast

Auxin is a plant growth hormone that plays a role in many aspects of growth and development, and its signalling have been also highly associated to plant stress responses ^48,49^. Several auxin signalling-related proteins were up accumulated in APF after infection (Table 1): PINOID (a RSTK involved in the regulation of auxin signalling, being a positive regulator of cellular auxin efflux ^48^); N-MYC DOWNREGULATED-LIKE2 (NDL2) protein (involved in auxin signalling through the positive regulation of auxin transporter protein 1 (AUX1) a carrier protein responsible for proton-driven auxin influx ^50^); and NEDD8 (neural precursor cell expressed developmentally down-regulated 8), a post-translational modifiers of cullin proteins ^51^. Cullin1 is one of the proteins that forms the “SKP1 cullin F-box” (SCF) ubiquitin ligase complex, that is involved in auxin signalling pathway ^52^. Auxin homeostasis could be perturbed by stress-induced changes that affects the auxin efflux carriers and that modify the apoplastic pH disturbing the auxin uptake and distribution ^53,54^. Indeed, in plant-pathogen interactions, auxin was shown to play a dual role both in regulating plant defence and in bacterial virulence ^55^. It was shown that some pathogens may promote host auxin accumulation which leads to the suppression of salicylic acid (SA)-mediated host defences. On the other side, host auxin accumulation also modulated expression of particular pathogen virulence genes ^55^. Despite the fact that none of the *P. viticola* effectors that we have found to be present in the APF was previously related to host auxin metabolism modulation, we may hypothesize that the modulation of auxin-related host proteins detected may reflect pathogen induced reprogramming. In fact, *P. viticola* being a biotroph, modulation of auxin metabolism leading to a depletion of SA-host defenses could be an infection strategy.

#### 3.1.4 Apoplastic ROS signalling in grapevine immunity

ROS play an important role in pathogen resistance by directly strengthening host cell walls via cross-linking of glycoproteins, promoting lipid peroxidation and activation of ROS-signalling networks ^56–58^. We have previously shown that ROS accumulation occurs in ‘Regent’ at 6hpi with *P. viticola* ^13,59^. In the APF proteome, proteins involved in ROS production, like peroxidase 4-like and purple acid phosphatase (PAP), are up-accumulated in response to the infection (Table 1; Fig.5), which is in accordance with our previous results. In Arabidopsis, PAP5 is induced in the early stages (6hpi) of *Pseudomonas syringae* infection and is involved in ROS generation ^60^. Apoplastic ROS are very important in plant development and responses to stress conditions, being involved in the activation of signal transduction from extracellular spaces to the cell interior and may directly eliminate invading pathogens (reviewed in ^61–63^). Small amounts of ROS lead to expression of stress-responsible genes as a plant resistance mechanism. However, high levels of ROS during a long period of time could culminate in damage of plant molecules and consequently in cell death. A strict regulation of ROS levels is thus important to induce a resistance response by the plant without promoting cell injury. In ‘Regent’ APF after infection with *P. viticola*, an up accumulation of xanthine dehydrogenase 1 (XDH1)-like isoform X1 was identified (Table 1; Fig.5). In Arabidopsis, this protein has dual and opposing roles in the ROS metabolism, contributing to H_2_O_2_ production in epidermal cells to fight pathogen haustoria and producing uric acid to scavenge chloroplast H_2_O_2_ in mesophyll cells to minimize oxidative damage ^64^. In grapevine, the role of this protein was not been elucidated yet. However, in our results, an up accumulation of this ROS-related protein in ‘Regent’ APF after infection was detected, suggesting a possible involvement of XDH1 in plant defence mechanism through ROS metabolism regulation.

In whole leaf proteome, an up accumulation of ROS-related proteins such as peroxisome biogenesis protein 19-2-like (PEX19-2) ^13,65,66^, protein TIC 62 (TIC62) ^67^ and succinate dehydrogenase assembly factor 4 (SDHAF4) ^68^ was observed at the same time that proteins like peroxidases and polyphenol oxidase ^69^ were less accumulated after infection (Table 2; Fig.5). Also, several heat shock proteins with different abundance levels in infected leaves, when compared to non-infected leaves, were identified (Table 2; ^70^). As we already mentioned, the regulation of ROS metabolism during plant-pathogen interaction presents a complex dynamic. ROS are generated to overcome the infection but at the same time a tight regulation of high ROS levels is needed to protect the plant from oxidative stress (reviewed in ^61–63^). This regulation is clearly evident in ‘Regent’ whole leaf and APF through the presence of several ROS-related proteins at different abundance levels, highlighting the importance of ROS in the ‘Regent’ defence mechanism against *P. viticola* as previously reported ^13,59^.

#### 3.1.5 Calcium related signalling is prevalent in the APF at 6hpi

In grapevine-*P. viticola* interaction, the increase in calcium leaf concentration in response to infection was already reported ^71^. In ‘Regent’ APF proteome we have found an up accumulation of a putative calcium-transporting ATPase 13 (Ca^2+^-ATPase), (Table 1; Fig.5). Ca^2+^-ATPases play critical roles in sensing calcium fluctuations and relaying downstream signals by activating definitive targets, thus modulating corresponding metabolic pathways (reviewed in ^72^). We have also detected an up accumulation of PINOID (Table 1). In Arabidopsis, PINOID activity is regulated by several calcium binding proteins, like a calmodulin-like protein, and the binding of these proteins to PINOID is enhanced by calcium ^73^.

In *P. viticola* proteome, we have detected a Ca^2+^/calmodulin-dependent protein kinase (CaMK) that responds to high levels of calcium ^74^, (Table 3; Fig.5). In pathogens, CaMKs are involved in several pathogenicity-related cellular mechanism. For example, in *C. albicans*, CaMKs functions in cell wall integrity and cellular redox regulation ^75^; in *Neurospora*, a genus of Ascomycete fungi, CaMKs are related to growth and development of the pathogens ^76^; and in *M. oryzae*, conidial germination and appressorial formation were delayed and virulence was attenuated in mutants of a CaMK ^77^. However, in *P. viticola*, up to our knowledge, there is no information about the role of these type of kinases in oomycete development and/or pathogenicity.

At the same time, in ‘Regent’ whole leaf proteome, calcium-related proteins, like calcium sensing receptor ^78^ and CDGSH iron-sulfur domain-containing protein NEET ^79^, are less accumulated after infection (Table 2).

These results suggest that, at such an early stage of the infection such as 6hpi, this calcium-associated response and regulation, as consequence of high levels of calcium in the infection site, is only taking place in the apoplast, which is the first contact point between plant and pathogen. As such, this regulation of the abundance of proteins associated with calcium metabolism in the leaf tissue is not evident.

#### 3.1.6 Activation of enzymes to disrupt oomycete structures

One of the plant defence mechanisms during pathogen infection is the secretion of proteins involved in the degradation of pathogen structures to inhibit its growth and thus stop its proliferation. Oomycete cell walls consist mainly of β-1,3-glucans, β-1,6-glucans and cellulose rather than chitin, essential constituent of fungal cell walls ^80^.

In grapevine APF after *P. viticola* infection an up accumulation of two glucan endo-1,3-beta-D-glucosidases and two GDSL esterase/lipases (GELPs) was observed (Table 1; Fig.4). The first are involved in the degradation of the polysaccharides of the pathogen cell wall and the GELPs possess lipase and antimicrobial activities that directly disrupt pathogen spore integrity ^81^. In contrast, in whole leaf proteome, proteins like endo-1,3;1,4-beta-D-glucanase-like, endo-1,3;1,4-beta-D-glucanase-like isoform X3 and glucan endo-1,3-beta-glucosidase, are less abundant after infection when compared to the non-infected leaves (Table 2; Fig.4). These results suggest that a demobilization of these proteins from the inside of the cell to the APF might be occurring in response to *P. viticola* infection. The accumulation of these proteins in ‘Regent’ APF leads to a disruption of oomycete structures as defence mechanism to inhibit the infection progress.

Moreover, a protein with antifungal activity, profilin 1, was found to be accumulated in ‘Regent’ whole leaf after *P. viticola* infection (Table 2). In Arabidopsis, this protein showed significant intracellular accumulation and cell-binding affinity for fungal cells, being capable to penetrate the fungal cell wall and membrane and act as inhibitor of fungal growth through ROS generation ^82^.

#### 3.1.7 The importance of protein trafficking during grapevine-*P. viticola* interaction

During plant-pathogen interaction, protein trafficking is very important for plant cells to quickly respond to pathogen infection. This trafficking occurs through the secretory and endocytic pathways that involves a complex set of proteins associated to vesicle formation, transport, docking, and fusion with the respective target membrane (reviewed in ^83^). Clathrin-mediated endocytosis is the best-known mechanism of endocytosis in plants and involves the generation of small vesicles surrounded by a coat of clathrin and other associated proteins. During interaction of ‘Regent’ with the oomycete *P. viticola*, we observed an accumulation of clathrin assembly protein in the whole leaf proteome at the first hours of infection (Table 2). Indeed, the increase of proteins involved in the generation of trafficking vesicles, like clathrin assembly protein, reinforces our hypothesis of a relocation of specific proteins within the cell and to the APF as a plant defence mechanism. We have already mentioned the presence of the protein unc-13 homolog in the APF, which is responsible for the translocation of H^+^-ATPases to the plasma membrane ^44^ and we have also proposed the translocation of glucanases from the inside of the cell to the APF in response to the infection. These results highlight the massive molecular reprogramming that occurs when a plant is exposed to an environmental stress like pathogen infection.

Even for the pathogen, the molecular trafficking that occurs within the cells is very important for its growth and pathogenicity. In *P. viticola*, we have identified the small GTP-binding Rab28 (Ras homologue from brain), (Table 3). Rab proteins, which constitute the largest family of monomeric GTPases, are small proteins involved in many biological processes (reviewed in ^84^). Members of this family participate in cell regulation, growth, morphogenesis, cell division, and virulence. Also, they are known as master regulators of intracellular bidirectional vesicle transport and, consequently, they localize in ER, vesicles, and multivesicular bodies, as well as in early and late endosomes ^85^. Rabs have been implicated in regulating vesicle motility through interaction with both microtubules and actin filaments of the cytoskeleton ^86^. In fungi, Rab participates in the secretion of metabolites and lytic enzymes and, in *Fusarium graminearum*, Rab GTPases are essential for membrane trafficking-dependent growth and pathogenicity ^87^. However, in *P. viticola* the specific function of Rab proteins was not yet elucidated.

#### 3.1.8 Host and pathogen proteases as hubs during the interaction

Proteases are enzymes that catalyse the breakdown of proteins into smaller polypeptides or single amino acids and play important roles in numerous biochemical processes. Pathogens produce a variety of proteases to degrade host tissue or to disrupt or modify host defence to create suitable conditions for successful colonization (reviewed in ^88,89^). In *P. viticola*, a calpain-like protease was detected (Table 3) and a possible role in pathogenicity is suggested as previous studies in *M. oryzae* point out that calpains play multiple roles in conidiation, sexual reproduction, cell wall integrity and pathogenicity ^90^. In *Saccharomyces cerevisiae* a calpain is also required for alkaline adaptation and sporulation ^91^. Moreover, two trypsins and serine proteases involved in pathogenicity ^28^ were identified in *P. viticola* proteome during grapevine infection (Table 3).

Host subtilisin-like protease SBT5.3, was found more accumulated in the APF after infection. This accumulation is in accordance with the previously published studies on subtilisin-like proteases in grapevine-*P. viticola* interaction. A 10-fold change in SBT5.3 gene expression was observed also at 6hpi together with the increase of expression of several subtilase genes in ‘Regent’ ^15^. Previously, it has also been hypothesized that subtilases play a crucial role in the establishment of the incompatible interaction between ‘Regent’ and *P. viticola*. In Arabidopsis, the subtilase SBT3.3 was shown to accumulate in the extracellular matrix after infection and to initiate a downstream immune signalling process ^92^.

Our results highlight that both host and pathogen proteases are essential on the first contact hours and might play an important role in pathogen recognition and in the overcome of host defences.

#### 3.1.9 *P. viticola* proteome at 6hpi reflects actively regulated processes leading to infection development

In *P. viticola*, several kinases are putatively involved in the infection mechanism were also found in the APF at 6hpi (Table 3). We have detected, in the *P. viticola* proteome, a AGC kinase (cAMP-dependent, cGMP-dependent and protein kinase C), (Table 3; Fig.5). This protein family embraces a collection of Ser/Thr kinases that mediate a large number of cellular processes, such as cell growth, response to environmental stresses, and host immunity ^93–96^. A study using kinase inhibitors showed that kinase C protein plays a key role in the signal transduction mechanisms during maintenance of motility of the zoospores, essential during their migration to the stomata ^97^. This evidence suggested that AGC kinases might be important pathogenic factors during pathogen infection, leading us to hypothesize that the *P. viticola* AGC kinase might be also relevant for the pathogenicity of this oomycete.

We have also detected a CDK (cyclin-dependent kinase) in *P. viticola* proteome that might be participating in pathogen infection mechanisms through cell polarization for the germ tube formation (Table 3; Fig.5). Indeed, several CDK members have been reported to be pathogenicity-related. In phytopathogenic fungus *U. maydis*, a CDK that is essential for growth and maintenance of cell polarity in this pathogen was identified. *cdk5ts* mutants showed to be drastically less virulent, probably because of the involvement of this protein in the induction of the polar growth required for the infection process ^98^. In *P. viticola*, cell polarity guides the emergence and the growth of the germ tube during the infection mechanism, namely the penetration into the stomatal cavity ^99,100^.

A MAPK (Mitogen-Activated Protein Kinase) has also been identified in the *P. viticola* proteome during the interaction with ‘Regent’ leaves (Table 3; Fig.5). MAPK cascades are very important in numerous cellular mechanisms in pathogens, regulating infection-related morphogenesis, cell wall remodelling, and high osmolarity stress response (reviewed in ^101,102^). For example, in *Peronophythora litchii*, the oomycete pathogen causing litchi downy blight disease, a mutation in PlMAPK10, a MAPKP, led to a reduced mycelial growth rate, less sporulation and weakened pathogenicity, indicating an important function of MAPK signal pathway in oomycete pathogenicity ^103^. In *Phytophthora sojae*, another oomycete, the authors showed that the MAPK PsSAK1 controls zoospore development and that it is necessary for pathogenicity ^104^. In *M. oryzae*, initiation of appressorium formation is controlled by the cyclic AMP-protein kinase A pathway and the appressorium development and invasive growth is regulated by the Pmk1 MAPK pathway ^105,106^.

Tyrosine kinase-like (TKL) was also identified in *P. viticola* proteome after 6 hours of infection (Table 3; Fig.5). The TKL family is present in most eukaryotes and participates in many biological processes, however, information about its role in oomycetes is still scarce. In *P. guiyangense*, TKLs were up-regulated at early infection stages and silencing of TKLs led to reduced mycelia growth, zoospore production and alteration of stress responses. Also, silencing of TKLs resulted in a reduced virulence of *P. guiyangense* ^107^, suggesting a key role of these kinases in pathogen infection strategies. In *P. infestans* kinome prediction 139 TKLs were identified, however their function is still unknown ^108^.

Tetratricopeptide repeat (TPR)- and SEL1-containing proteins were also identified in *P. viticola* proteome (Table 3). These have been reported to be directly related to virulence-associated functions in bacterial pathogens ^109–111^. In *Francisella tularensis*, a bacterial pathogen, a TPR-containing protein is a membrane-associated protein that is required for intracellular replication of the microbe, *in vivo* virulence, and heat stress tolerance ^112^. In *P. infestans*, TPR was predicted as one of the most common domains within proteins ^113^. However, up to our knowledge, the role of TPR-containing proteins in oomycete/fungus pathogenicity is still unknown. Regarding SEL1-containing proteins, in *Candida albicans*, a human fungal pathogen, these proteins are capable of shaping host immune response and the severity of fungal systemic infection and were suggested as a novel fungus-derived pattern-associated molecular pattern (PAMP) ^114^. However, up to our knowledge, there is no information about pathogen Sel1-containing proteins in plant-oomycete interaction.

Growth-related proteins were also detected in the *P. viticola* proteome sequencing (Table 3; Fig.4). These types of proteins are essential for the development of the pathogen structures during infection and besides they are not directly related to pathogenicity, their presence is an indicator of the pathogen growth and thus the progression of the infection in the host. In *P. viticola*, we have identified the SKN1 protein that is required for synthesis of the β-glucans, the major components of oomycete cell walls ^115^. We have also identified the cell division cycle 5 (Cdc5) protein, a highly conserved nucleic acid binding protein among eukaryotes that plays critical roles in development, however, in pathogens this protein is still poorly characterized ^116^.

## 4. Conclusions

Plants have developed several defence mechanisms to rapidly respond to pathogen attack. A fast recognition of the pathogen structures and activation of a defence response is primordial for the establishment of the incompatible interaction. Apoplast dynamics becomes an essential part of the battle between plants and pathogens. Here, we reveal for the first time the communication between grapevine extracellular and intracellular spaces at the first hours of interaction with *P. viticola* as well as the pathogen strategies to overcome grapevine defence. Our results highlight several defence mechanisms that are first activated and modulated in grapevine apoplast during infection, namely leading to plant cell wall plasticity to prevent disruption and disturbance of oomycete structures. Several proteins were also identified in the whole leaf proteome of ‘Regent’ that are closely related with the mechanisms involved in the apoplast modulation during infection, highlighting a tight communication between the APF and cell interior. Moreover, we have shown that *P. viticola* proteome is enriched in virulence-related proteins as a strategy to defeat the plant defence response and in growth-related proteins to develop the infection structures of the *P. viticola*.

The analysis of the plant and oomycete proteins involved in the first hours of the interaction between grapevine and *P. viticola* revealed that both sides are modulating their offense and defence strategies very early on. This reinforces the hypothesis that host-pathogen cross-talk in the first hours of interaction is highly dynamic and processes such as ETI, ETS, PTI and PTS may occur simultaneously.

## 5. Materials and Methods

### 5.1 Plant material and inoculation experiments

The tolerant *V. vinifera* cv. ‘Regent’ (VIVC number 4572) was used in this study. Wood cuttings from ‘Regent’ were obtained at the Portuguese Grapevine Germplasm Bank (^117^; INIAV – Dois Portos, Portugal) and grown in 2.5 L pots in universal substrate under controlled conditions in a climate chamber at natural day/night rhythm, relative humidity 60% and a photosynthetic photon flux density of 300 μmol m^-2^ s^-1^.

For plant inoculation, downy mildew symptomatic leaves were harvested at the Portuguese Grapevine Germplasm Bank, sprayed with water and incubated overnight at 22°C in the dark to enhance sporulation. Next day, *P. viticola* sporangia were collected and their vitality was checked by microscopy. Inocula was propagated in the laboratory using the susceptible Müller-Thurgau.

For the experimental assay, ‘Regent’ abaxial leaf surface was sprayed with an inoculum solution containing 3.5×10^−5^ sporangia/mL. Mock-inoculations with water were also made and used as control. After inoculation, plants were kept in a greenhouse under high humidity conditions and 25°C. The third to fifth fully expanded leaves beneath the shoot apex were harvest at 6 hours post-inoculation for apoplastic fluid extraction.

### 5.2 Apoplastic fluid extraction

Apoplastic fluid and total soluble protein extraction was performed as described in ^3^. Briefly, 5 volumes of 0.1 M ammonium acetate in methanol were used to precipitate the proteins overnight at -20°C. Samples were centrifuged at 4000 *g*, during 30 min at -10°C. Recovered pellets were washed (once with 0.1 M ammonium acetate in 100% methanol, twice with 80% (v/v) acetone and twice with 70% (v/v) ethanol), dried and resuspended in 0.03 M Tris-HCl buffer (pH 8.8) solution containing, 7 M urea, 2 M thiourea, 4% (w/v) CHAPS ^3^. For cytoplasmic content contamination control, malate dehydrogenase assay was used according to ^3^.

### 5.3 MS-Based protein identification

Separation and MS-based identification of proteins was performed as described in ^3^. For identification of *V. vinifera* and *P. viticola* proteins, the genome assembly of *Vitis*12X database (GCF-000003745.3; 41 208 sequences, July 2009) and *Plasmopara viticola* genome database (INRA-PV221 isolate; 15 960 sequences, April 2018), respectively, were used via Mascot Daemon (v.2.6.0. Matrix Science), imported to Progenesis QIP and matched to peptide spectra. The Mascot research parameters were: a peptide tolerance of 20 ppm, a fragment mass tolerance of 0.3 Da, carbamidomethylation of cysteine as fixed modification and oxidation of methionine, N-terminal protein acetylation and tryptophan to kynurenine as variable modifications. Only the proteins identified with a significance MASCOT-calculated threshold P-value < 0.05, at least two significant peptides per proteins and one unique peptide per proteins were accepted.

For identification of *P. viticola* proteins present in apoplast of ‘Regent’ leaves inoculated with the oomycete, only the sequenced proteins that fulfilled the following two criteria at the same time were considered: be present in at least 2 of the biological replicates in the inoculated samples; and present in only 1 biological replicate or totally absent in control samples. Functional information about *P. viticola* proteins was obtained from *P. viticola* genome database. Protein secretion and effector function were predicted using TargetP 2.0 (https://services.healthtech.dtu.dk/service.php?TargetP-2.0; ^118^) and EffectorP (http://effectorp.csiro.au/; ^119^), respectively.

### 5.4 Whole leaf proteome data

For ‘Regent’ whole leaf proteome analysis, 6hpi with *P*.*viticola*, an already published dataset deposited on the ProteomeXchange Consortium via the PRIDE partner repository with the identifier PXD021613 was used ^13^.

### 5.5 Statistical analysis of APF and whole leaf proteomes

Principal Component Analysis (PCA) of ‘Regent’ APF proteome at 6hpi with *P. viticola* was performed using the program MetaboAnalyst 5.0 (http://www.metaboanalyst.ca/, ^120^).

For the statistical analysis and consequent identification of differentially accumulated proteins, both whole leaf and APF proteomes of ‘Regent’ leaves inoculated with *P. viticola* (6hpi) were matched to the respective control samples, allowing a comparative analysis between datasets. This was done by applying Rank Products (RP), a powerful statistical test designed for identifying differentially expressed genes in microarrays experiments ^121^, nevertheless, it also provides a simple and straightforward tool to determine the significance of observed changes in other omics data. RP procedure makes weak assumptions about the data and provides a strong performance with very small data sets, allowing for the control of the false discovery rate (FDR) and familywise error rate (probability of a type I error) in the multiple testing situation.

### 5.5 Subcellular location prediction of APF DAPs

For the APF DAPs from grapevine, a subcellular localization prediction was performed using SignalP 5.0, TargetP 2.0 and SecretomeP 2.0 servers (http://www.cbs.dtu.dk/services/, ^122–124^), ApoplastP (http://apoplastp.csiro.au/, ^125^), BUSCA (http://busca.biocomp.unibo.it/, ^126^), PredSL (http://aias.biol.uoa.gr/PredSL/, ^127^), Mercator (https://www.plabipd.de/portal/mercator-sequence-annotation, ^128^) and Blast2GO (version 5.2.5, https://www.blast2go.com/, ^129^). The default parameters were used for all the programs.

Based on the subcellular localization prediction, the APF proteins were grouped in 4 different classes according to the following criteria (as described in ^3^): (1) proteins with a predicted signal peptide (SP) by SignalP (Class I; (2) proteins predicted to be secreted through classical secretory pathways but, by other software than SignalP 5.0 (Class II); (3) proteins predicted to be secreted by unconventional secretory pathways (USP) based on SecretomeP (Class III), and proteins with no predicted secretion (Class IV). Only the proteins belonging to the Class I, II and III were considered for further functional analysis.

### 5.6 APF and whole leaf proteome data functional analysis

Functional annotation based on Gene Ontology annotation (Biological Process) using Blast2GO software was performed.

## Data Availability

The mass spectrometry proteomics data have been deposited to the ProteomeXchange Consortium via the PRIDE partner repository with the dataset identifier PXD030508 and 10.6019/PXD030508.

## Acknowledgments

Portuguese Foundation for Science and Technology (FCT, Portugal) funded Joana Figueiredo fellowship (SFRH/BD/137066/2018). FCT funded the Research Units and project: BioISI (UIDB/04046/2020 and UIDP/04046/2020), LEAF (UID/AGR/04129/2019) and CEAUL (UIDB/00006/2020), and the project PTDC/BIA-BQM/28539/2017. We thank Dr. Ana Rita Cavaco, Dr. Marisa Maia and Dr. Maria do Céu Silva for the support in the inoculation assay and APF extraction protocol.

## Conflicts of Interest

The authors declare no conflict of interest. The funders had no role in the design of the study; in the collection, analyses, or interpretation of data; in the writing of the manuscript, or in the decision to publish the results.

## Author Contributions

J.F. and A.F. conceived the study; J.F., R.B.S, L.G.G. and A.F. were responsible for the plant material and performed the APF extraction method; J.F. prepared the samples for nanoLS-MS/MS analysis; C.C.L. and J.R. performed the proteome profiling by nanoLC-MS/MS; L.S. performed the statistical analysis; J.F. and A.F. analysed the data and wrote the manuscript. All authors have read and agreed to the published version of the manuscript.

## Notes

### Competing Interest Statement

The authors have declared no competing interest.

